# GPR4 knockout attenuates intestinal inflammation and forestalls the development of colitis-associated colorectal cancer in murine models

**DOI:** 10.1101/2023.03.13.532341

**Authors:** Mona A. Marie, Edward J. Sanderlin, Alexander Hoffman, Kylie D. Cashwell, Swati Satturwar, Heng Hong, Ying Sun, Li V. Yang

**Author notes:** Corresponding author (LVY).

## Abstract

GPR4 is a proton-sensing G protein-coupled receptor highly expressed in vascular endothelial cells and has been shown to potentiate intestinal inflammation in murine colitis models. Herein, we evaluated the proinflammatory role of GPR4 in the development of colitis-associated colorectal cancer (CAC) using the dextran sulfate sodium (DSS) and azoxymethane (AOM) mouse models in wild-type and GPR4 knockout mice. We found GPR4 contributed to chronic intestinal inflammation and heightened DSS/AOM-induced intestinal tumor burden. Tumor blood vessel density was markedly reduced in mice deficient in GPR4 which correlated with increased tumor necrosis and reduced tumor cell proliferation. These data demonstrate GPR4 ablation alleviates intestinal inflammation and reduces tumor angiogenesis, development, and progression in the AOM/DSS mouse model.

**Author summary:** Inflammatory bowel disease (IBD), including ulcerative colitis and Crohn’s disease, is a debilitating condition with chronic inflammation in the digestive tract. Patients with IBD are at higher risk of developing colitis-associated colorectal cancer (CAC), compared with the general population. The etiology of IBD is not well understood, but both genetic and environmental factors have been implicated. In this study, we investigated the role of the pH-sensing GPR4 receptor in colitis and CAC using the DSS and AOM induced mouse models. GPR4 knockout alleviated intestinal inflammation, reduced tumor angiogenesis, and impeded CAC development. Our data suggest that inhibition of GPR4 may be explored as a potential therapeutic approach for IBD treatment and CAC prevention.

## 1. Introduction

Colitis-associated colorectal cancer (CAC) is driven by chronic and recurring mucosal inflammation observed in inflammatory bowel disease (IBD) patients [1]. Patients with IBD have an increased risk of developing CAC when compared to the general population [2]. In this context, colorectal cancer (CRC) can develop and progress through the chronic inflammation-dysplasia-carcinoma axis [1]. Interestingly, severe intestinal inflammation is associated with decreased colon luminal pH in IBD patients [3,4,5]. Tissue acidosis is characterized by reduced extracellular pH and is a hallmark of chronic inflammation and cancer. As such, solid tumors are characterized by an acidic tumor microenvironment [6,7,8]. Local tissue acidification of inflamed colonic tissues was also observed in a DSS-induced colitis mouse model [9]. Inflamed colon segments excised from DSS-treated mice were more acidic than non-inflamed colon segments as measured by the pH indicator, SNARF 4 F-5 (and 6) carboxylic acid [9].

A family of proton-sensing G protein-coupled receptors (GPCR) capable of sensing changes in extracellular pH have recently been implicated in regulating intestinal inflammation. The pH-sensing GPCR family includes GPR4, GPR65, and GPR68 of which are activated by protonation of histidine residues on the receptor extracellular domains [7,10,11,12,13,14,15]. GPR4 is predominately expressed on vascular endothelial cells among other cell types [16,17,18,19,20,21,22]. Our group has demonstrated that GPR4 activation by acidic microenvironment augments endothelium adhesiveness and increases the expression of endothelial proinflammatory molecules, such as IL-8, CXCL2, COX-2, VCAM-1 and E-selectin [17,23]. Moreover, intestinal fibrosis is a serious complication in IBD [24,25]. GPR4 expression is positively correlated to fibrogenic gene expression in highly fibrotic intestine lesions obtained from Crohn’s disease (CD) patients [20]. In addition to proinflammatory, profibrotic and endothelial cell activation roles, GPR4 has been implicated in both physiological and pathological angiogenesis [19,20,26,27,28]. Studies demonstrated a role for GPR4 in promoting angiogenesis and tumor growth in the porous tissue implant and orthotopic tumor models [28]. GPR4 expression has been associated with increased angiogenesis in hepatocellular, head and neck, breast and colorectal cancers [27,28,29]. Increased GPR4 expression was observed in colorectal tumors compared to adjacent normal tissue and was associated with decreased overall survival in patients [30]. Therefore, GPR4 may contribute to tumorigenesis in the colon through reinforcing the inflammatory and angiogenic processes in IBD. Herein, using wild-type and GPR4 knockout mouse CAC models, we investigated the role of GPR4 in inflammation driven colorectal cancer.

## 2. Materials and Methods

### 2.1. Ethics statement

The mouse experiments were approved by the Institutional Animal Care and Use Committee of East Carolina University in accordance with the *Guide for the Care and Use of Laboratory Animals* (The National Academies Press).

### 2.2. Dextran sulfate sodium (DSS)-induced colitis mouse model

9 weeks old male and female wild-type (WT) and GPR4 knockout (KO) mice were used in the experiments. GPR4 deficient mice were generated as previously described and were backcrossed into the C57BL/6 background for 11 generations [19]. Animals were maintained under specific pathogen-free conditions and were free from *Helicobacter, Citrobacter rodentium*, and *norovirus*. Colitis was induced using 3% (w/v) colitis grade dextran sulfate sodium (DSS) with molecular weight 36,000-50,000 Da (Lot# Q1408, MP Biomedical, Solon, OH) within the drinking water of mice. The 3% DSS solution or water was provided to mice *ad libitum*, as previously described [31]. Briefly, mice were given 3% DSS for 4 cycles. Each cycle constituted 5 days of 3% DSS followed by 2 days of water. Following the fourth cycle, water was switched back to 3% DSS for 2 final days. Mouse body weight and clinical phenotype scores were measured each day [31].

### 2.3. Azoxymethane (AOM) and dextran sulfate sodium (DSS)-induced colitis associated colorectal cancer mouse model (CAC)

9 to 12 weeks old male and female WT and GPR4 KO mice were used in these experiments. Mice were backcrossed 13 generations into the C57BL/6 background. CAC mouse model was induced as previously described [31]. Briefly, a single *i*.*p*. injection (10 mg/kg) of azoxymethane (AOM, product# A5486, Sigma-Aldrich, Saint Louis, MO) followed by 4% (w/v) DSS (Lot# Q5229 and S0948, MP Biomedical, Solon, OH) in drinking water was used to induce colitis-associated colorectal cancer in mice. The mice were given 4% DSS in water for 3 cycles. Each cycle constituted 5 days of 4% DSS followed by 16 days of water. Following the third cycle, mice were given water for the remaining period (37 days) to the endpoint on the 14^th^ week (100 days).

### 2.4. Mouse clinical phenotype scoring

Assessment of colitis severity in WT and GPR4 KO mice treated with 3% DSS or AOM/ 4% DSS was performed as previously described [31]. Colitis severity was determined using the clinical parameters of body weight loss and fecal score [16]. Disease activity index represented by body weight loss percentage, fecal score, colon shortening, and mensenteric lymph node expansion, was measured to assess inflammation. Feces was collected from mice and assessed for presence of blood and consistency. Fecal scoring system consisted of the following: 0= normal, dry, firm pellet; 1= formed soft pellet with negative hemoccult test, 2= formed soft pellet with positive hemoccult test; 3= formed soft pellet with visual blood; 4= liquid diarrhea with visual blood; 5= bloody mucus and no colonic fecal content upon necropsy. Presence of micro blood content was measured using the Hemoccult Single Slides screening test (Beckman Coulter, Brea, CA).

### 2.5. Mouse tissue collection, evaluation, and processing

At endpoint, mice were euthanized followed by necropsy, colon, and mesenteric lymph nodes were collected, evaluated and processed, as previously described [16,31]. Colon length was measured from the ileocecal junction to anus. Then colon was separated from cecum and phosphate buffer saline (PBS) was used to clear it of fecal content then it was opened along the anti-mesenteric border. Colon tissue was fixed with 10% buffered formalin and cut evenly into distal, mid, and proximal sections for histologic evaluation. The mesenteric lymph node and tumor volumes were assessed with a caliper using the formula (length × width^2^) π/6 using a dissecting scope for visualization then fixed with 10% buffered formalin and collected for histological analysis.

### 2.6. Histopathological analysis

Five µm sections of distal, middle, and proximal colon tissue segments were obtained from WT and GPR4 KO mice and stained with hematoxylin and eosin (H&E) for histological analysis. Sample identification was concealed during histopathological analysis for unbiased evaluation. Board certified medical pathologists evaluated colitis histopathological features including inflammation, crypt damage, edema, architectural distortion, and leukocyte infiltration in a blinded manner as previously described [16,32,33]. Each parameter was scored and multiplied by a factor corresponding to total tissue percentage of disease involvement. Additional scoring criteria of colonic fibrosis were evaluated as previously described [34]. Briefly, colon segments of WT and GPR4 KO mice treated with DSS were stained with picrosirius red for fibrosis analysis and graded in a blinded manner for pathological fibrosis. Severity of fibrosis was included as one of the parameters of the histopathological score. WT and GPR4 KO AOM/DSS treated mouse intestinal tumors in the colons were assessed to differentiate dysplasia and adenocarcinoma lesions.

### 2.7. Tumor necrotic area quantification

Hematoxylin and Eosin (H&E) stain was performed on formalin fixed, paraffin embedded tumor sections to assess necrosis in WT and GPR4 KO AOM/DSS mouse colons. Percent of necrotic area per field of view (FOV) was measured using ImageJ software in a blinded manner. Briefly, images were taken for each tumor to capture the total tumor area using a microscope (Axio Imager M2, Zeiss Inc.). Percent of necrosis per FOV was calculated using the following equation: % necrotic area = (sum of necrotic areas / total area of FOV) × 100.

### 2.8. Immunohistochemistry

Colon tissues were embedded in paraffin and serial five µm sections were evaluated for immunohistochemical analysis as previously described [16]. Briefly, antigen retrieval was performed of colon sections followed by endogenous peroxidase blocking. Endogenous biotin, biotin receptors, and avidin binding sites were blocked using Avidin/ Biotin blocking kit (Vector laboratories, California, CA) followed by normal serum blocking and stained with anti-Green Florescence Protein (GFP) (Abcam, ab6673, Cambridge, MA), anti-TNF-α (Beijing Biosynthesis Biotechnology Co., Ltd., #BS-2081R, Beijing, China), anti-Cleaved Caspase-3 (Cell Signaling Technology, #9664S, Danvers, MA), anti-CD31 (Cell Signaling Technology, #77699S, Danvers, MA), and anti-Ki67 (Abcam, #ab15580, Waltham, MA) primary antibodies. The IHC detection system VECTASTAIN® Elite ABC-HRP Kit, Peroxidase (Rabbit IgG) (Vector laboratories, California, CA), and VECTASTAIN® Elite ABC-HRP Kit, Peroxidase (Goat IgG) (Vector laboratories, California, CA), were used. Following addition of secondary antibody, DAB (3,3’ – diaminobenzidine) incubation was performed for HRP detection. Pictures were taken using the Zeiss Axio Imager M2 microscope.

### 2.9. Microvessel density quantification

CD31^+^ immunohistochhemistry stain was used as a marker for endothelial cells forming blood vessels in the WT and GPR4 KO AOM/DSS colon tumor sections. Image J (v.1.53k) software was used for manual quantification of individual blood vessels/ field of view (FOV). 5-8 images were captured per tumor and blood vessel numbers were averaged. Images were analyzed in a blinded manner. Data is represented as the averaged tumor blood vessel number / FOV for the two groups WT and GPR4 KO AOM/DSS.

### 2.10. Tumor proliferation quantification

Ki67 immunohistochhemistry stain was used as a proliferation marker in the WT and GPR4 KO AOM/DSS colon tumor sections. Fiji (v.1.53q) software was used for analysis, as previously described [35]. Briefly, 4-8 images were captured per tumor and analyzed in a blinded manner. Percent positive cells were calculated by dividing the number of Ki67 positive (brown) cells by the total number of cells (blue nuclei)/ FOV multiplied by 100. Then all FOV percentages were averaged to produce a percentage per tumor.

### 2.11. Real time Polymerase Chain Reaction (RT-PCR)

Real-time PCR was performed as previously described [16]. Crohn’s and ulcerative colitis cDNA arrays were purchased from Origene Technologies (catalog #CCRT102, Rockville, MD) and utilized in real-time PCR analysis using specific primer-probes for human GPR4, TNF-α, IFN-γ and β-actin [23,31]. The cDNA array patient information were described in a previous report as supplied by the vendor [16].

### 2.12. Statistical analysis

All statistical analysis was performed using the GraphPad Prism (v.9.4.1) software. The unpaired t-test, Mann-Whitney U test, and one-way ANOVA were used to compare differences between two groups (WT vs. GPR4 KO) or more than two groups, respectively. Linear regression was used to correlate gene expression. P < 0.05 is considered statistically significant.

## 3. Results

### 3.1. GPR4 potentiates intestinal inflammation in the chronic DSS-induced experimental colitis mouse model

We used WT and GPR4 KO mice to characterize the role of GPR4 in the chronic DSS-induced colitis mouse model [36,37]. The model consists of 4 cycles of intestinal inflammatory insults. Mouse bodyweight and fecal blood and diarrhea were analyzed as disease activity indicators (Fig. 1). WT-DSS mice began to lose between 12-15% body weight following cycle one whereas GPR4 KO-DSS mice lost between 5-10% body weight throughout the cycles (Fig. 1A). Fecal scores also indicated GPR4 KO-DSS mice were less clinically severe when compared to WT-DSS mice as GPR4 KO-DSS mice had reduced fecal blood and diahrea (Fig. 1B). Upon completion of all four cycles of the chronic DSS-induced colitis, macroscopic disease indicators were collected such as mesenteric lymph node (MLN) enlargement and colon length measurements. MLN volume change due to inflammation was not different at this point of the study between WT and GPR4 KO-DSS mice (Fig. 1C). The colon length, however, indicated GPR4 KO-DSS mice had less colon shortening when compared to WT-DSS mice (Fig. 1D). Collectively, these results indicate GPR4 potentiates disease severity in the chronic DSS-induced mouse model.

**Fig. 1.**
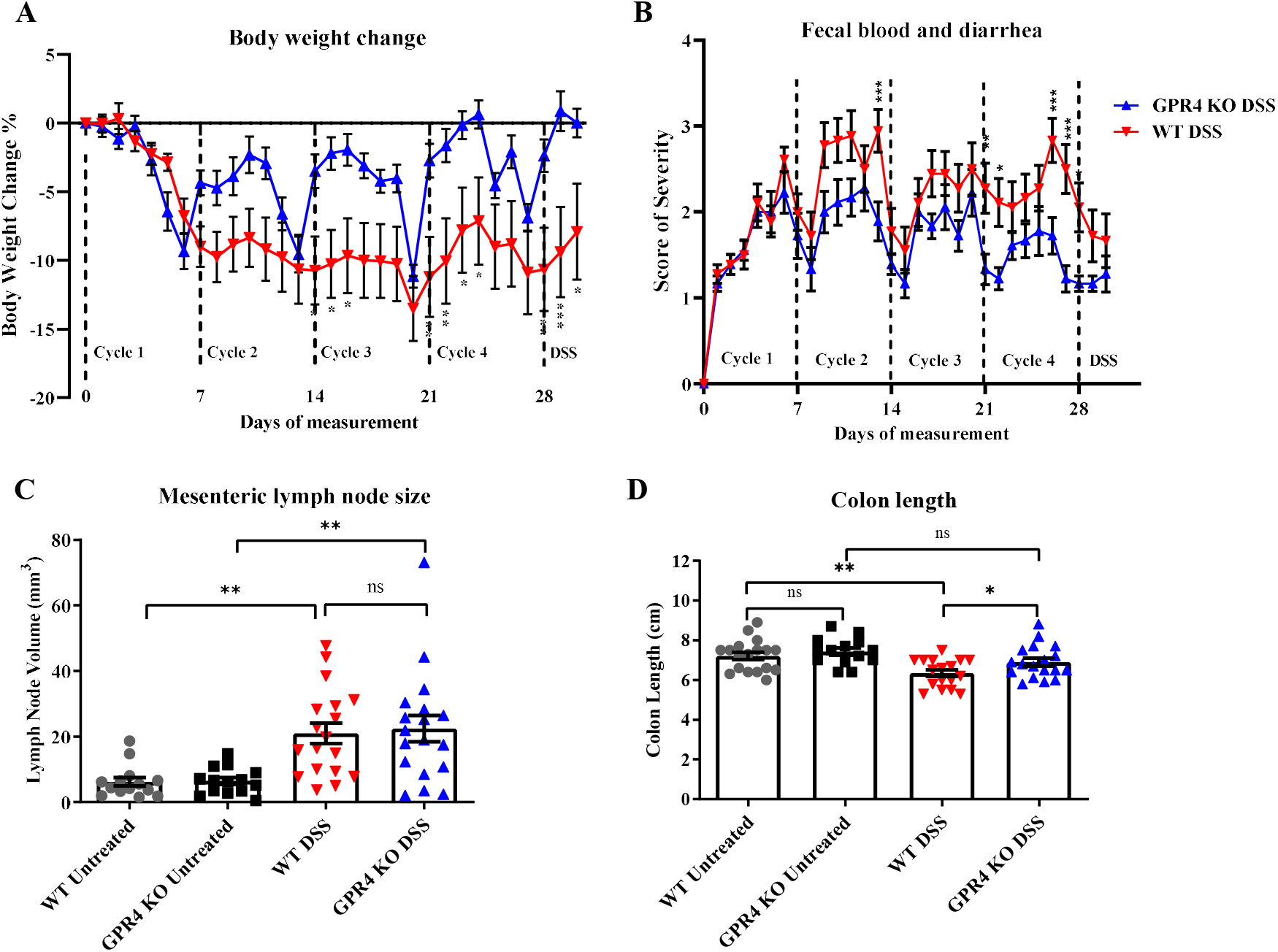
Disease activity indicators of chronic colitis induction in wild-type (WT) and GPR4 knockout (KO) mice. DSS-induced inflammation was assessed in WT-DSS and GPR4 KO-DSS mice. WT-DSS mice presented elevated inflammatory indicators compared to GPR4 KO-DSS mice. Clinical phenotypes of intestinal inflammation such as (A) body weight loss and (B) fecal blood and diarrhea were assessed. Macroscopic disease parameters such as (C) mesenteric lymph node expansion and (D) colon shortening were also recorded. Data are presented as the mean ± SEM and statistical significance was determined using the unpaired t-test between WT-DSS and GPR4 KO-DSS groups or one-way ANOVA test between the four groups, WT control, WT-DSS, GPR4 KO control and GPR4 KO-DSS. WT control (n=10), WT-DSS (n=13), GPR4 KO control (n=11), and GPR4 KO-DSS (n=13) mice were used for experiments. (*P < 0.05, **P < 0.01).

We then evaluated the degree of histopathology in the distal, middle, and proximal colon segments. Distinct parameters of colitis associated histopathology were assessed such as leukocyte infiltration, edema, crypt loss, and architectural distortion to obtain a score of severity. WT and GPR4 KO untreated water control mice displayed no observable histopathology (Fig. 2A). GPR4 KO-DSS mice had reduced histopathology when compared to WT-DSS mice in the colon segments (Fig. 2A-B).

**Fig. 2.**
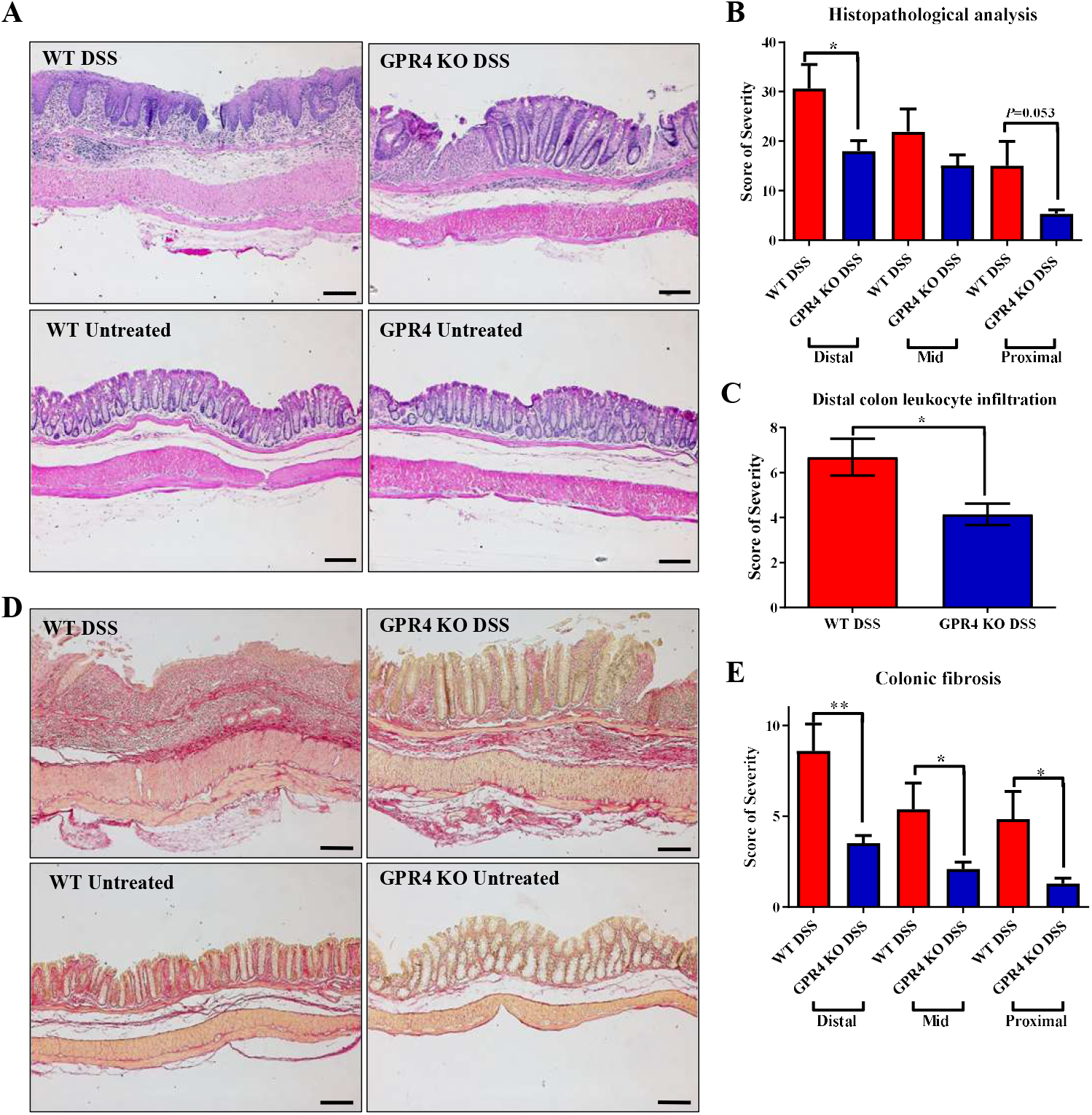
Histopathological analysis of proximal, middle, and distal colon in chronic colitis mice. Characteristic histopathological features of colitis were assessed to further characterize the degree of intestinal inflammation. WT-DSS mice presented elevated disease indicators compared to GPR4 KO-DSS mice. Representative H&E pictures were taken for (A) WT-DSS, GPR4 KO-DSS, WT control, GPR4 KO control mice, respectively. (B) Total histopathologic scores for WT-DSS and GPR4 KO-DSS proximal, middle, and distal colons. For fibrosis assessment, representative pictures of Picrosirius red stained tissue sections were taken for (D) WT-DSS, GPR4 KO-DSS, WT-control, and GPR4 KO control mice, respectively. Graphical representations of (B) total histopathological parameters, (C) Leukocyte infiltration assessment in the distal colons, and (E) colonic fibrosis are presented. WT control (n=10), WT-DSS (n=13), GPR4 KO control (n=11), and GPR4 KO-DSS (n=13) mouse tissues were used for histopathological analysis. Scale bar is 100µm. Data are presented as the mean ± SEM and statistical significance was determined using the unpaired t-test between WT-DSS and GPR4 KO-DSS groups. (*P < 0.05, **P < 0.01).

Moreover, leukocyte infiltration was reduced in the colon of GPR4 KO-DSS mice when compared to WT-DSS mice (Fig. 2C), which corroborates previous reports indicating GPR4 can increase leukocyte infiltration into inflamed intestinal tissues through upregulating endothelial cell adhesion molecules [16].

Another distinct histopathological consequence is fibrosis in chronically inflamed intestinal tissues. We observed heightened fibrotic development in mice with chronic DSS-induced colitis. The distal colon segment displayed the highest degree of fibrosis with a progressive reduction of severity from middle to proximal. A significant reduction in fibrosis was observed in the GPR4 KO-DSS mice in the distal, middle, and proximal colon segments when compared to WT-DSS mice (Fig. 2D-E).

### 3.2. GPR4 gene expression is positively correlated with TNF-α and INF-γ gene expression in the inflamed intestinal tissues

Previous studies have shown that GPR4 gene expression is upregulated in both ulcerative colitis (UC) and Crohn’s disease (CD) patient intestinal samples compared to normal intestinal tissue [16,18]. TNF-α and INF-γ are cytokines implicated in the inflammatory pathways of IBD [38]. Herein, we observed a positive correlation between the gene expression of GPR4 with both TNF-α (*P*< 0.0001 and *R*= 0.5966) and INF-γ (*P*< 0.0001 and *R*= 0.5634) in IBD patient intestinal samples (Fig. 3A-B). We also assessed the TNF-α protein expression in the distal colon sections of WT and GPR4 KO chronic DSS mice using immunhistochemistry. TNF-α protein expression was significantly reduced in the colon tissues of GPR4 KO chronic DSS mice compared to WT (Fig. 3C-D).

**Fig. 3.**
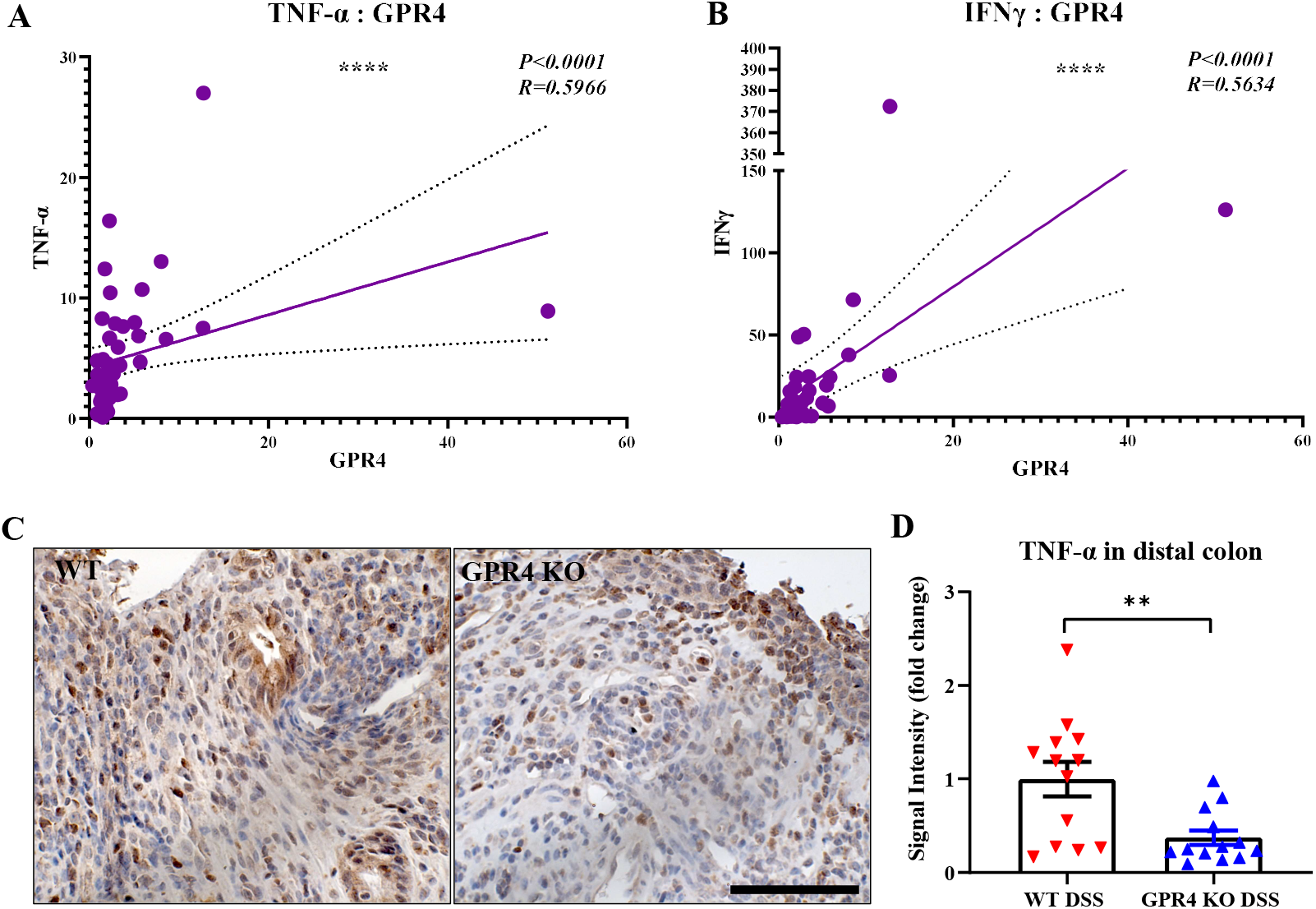
GPR4 effect on inflammatory mediators. GPR4 gene expression is correlated to inflammatory mediators TNF-α and INF-γ gene expression in colon tissue samples of IBD patients. A positive correlation was found between (A) GPR4 and TNF-α, and (B) GPR4 and INF-γ relative mRNA expression in IBD patients’ samples. Statistical significance was tested by Pearson correlation coefficient. TNF-α is downregulated in GPR4 KO-DSS mouse distal colons compared to WT-DSS. (C) Immunohistochemical representation of TNF-α in WT-DSS and GPR4 KO-DSS mouse distal colons, and (D) Semi-quantitative analysis of TNF-α protein expression in mouse distal colons of WT-DSS (n=13) and GPR4 KO-DSS (n=13). Unpaired t-test was used for statistical analysis. (**P < 0.01, ****P < 0.0001).

Genetic deletion of GPR4 reduces disease severity in CAC induced by AOM/DSS in mice To further study the role of GPR4 in the development of colitis associated colorectal cancer (CAC), we utilized the well established AOM/DSS murine model [39,40]. Disease severity indicators were measured during the experiment by monitoring body weight loss and fecal blood and diarrhea scores. WT AOM/DSS mice showed more significant body weight loss when compared to GPR4 KO AOM/DSS mice starting at the end of the first cycle, and throughout the second and third cycles. Both WT and GPR4 KO AOM/DSS mice reached an average body weight loss of ∼ 17% by day 9 during the first cycle, followed by a partial recovery in the body weight loss. This body weight recovery was more significant in the GPR4 KO mice than WT (Fig. 4A). The severity of fecal blood and diahrea scores was also significantly higher in WT AOM/DSS in comparison to GPR4 KO AOM/DSS mice (Fig. 4B). Upon the endpoint of the experiment (day 100), macroscopic disease indicators were evaluated such as final body weight, mesenteric lymph node expansion, and colon length shortening. GPR4 KO AOM/DSS mice showed a significant increase in body weight recovery compared to WT (Fig. 4C). Mesenteric lymph node volume of WT mice were larger than that of GPR4 KO mice, indicating a more severe intestinal inflammation in WT mice (Fig. 4D). No significant differences were observed in colon length between WT and GPR4 KO AOM/DSS-treated mice (Fig. 4E).

**Fig. 4.**
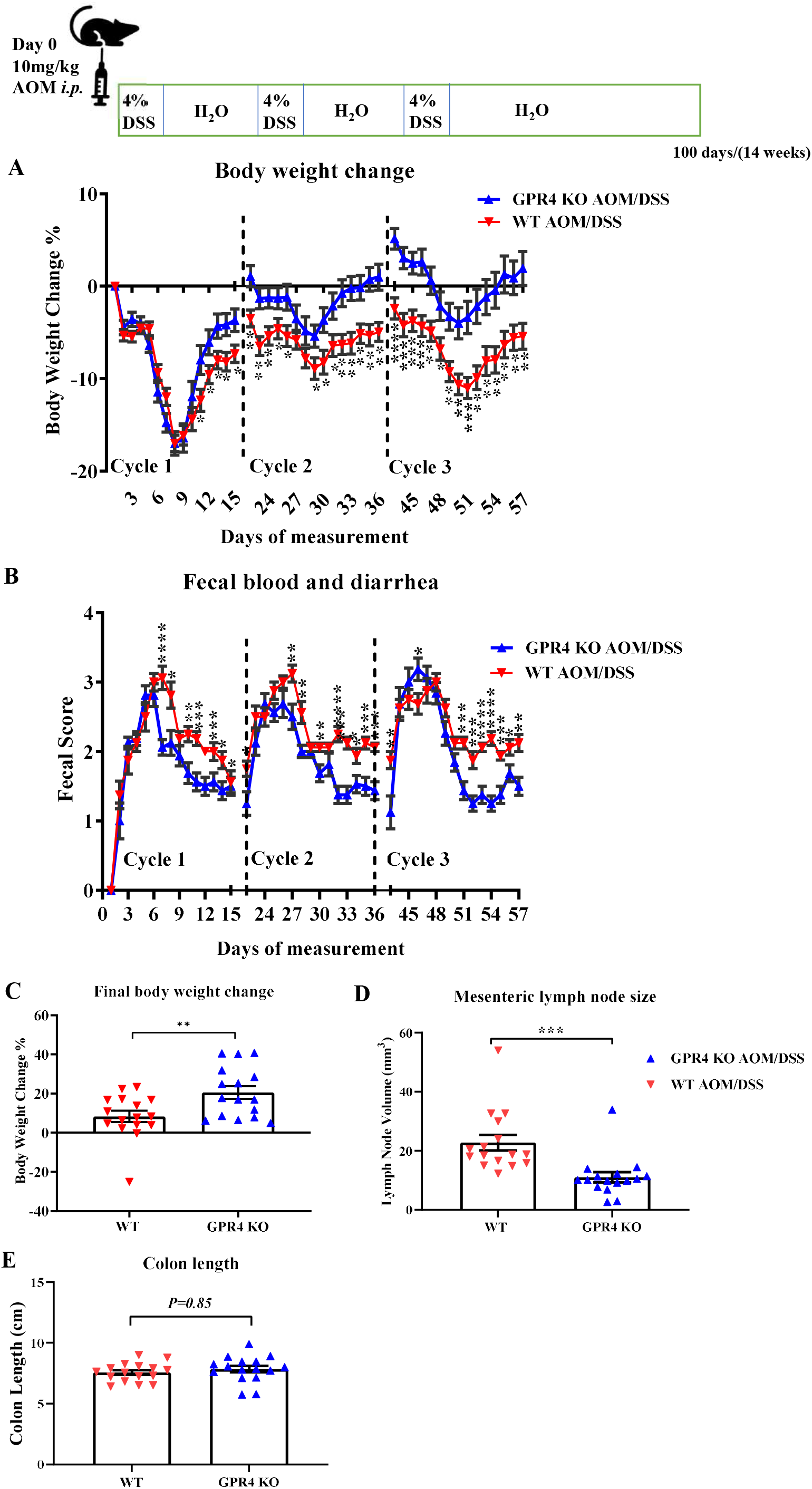
GPR4 knockout reduces intestinal inflammation in the colitis associated colorectal cancer (CAC) mouse model. WT AOM/DSS (n=16) and GPR4 KO AOM/DSS (n=16) mice were used for this analysis. WT AOM/DSS mice showed more servere body weight loss and fecal blood and diarrhea score throughout treatment cycles when compared to GPR4 KO AOM/DSS mice. (A) Body weight change percentage, and (B) fecal blood and diarrhea score. At the endpoint, disease parameters such as final body weight change, colon length, and menseteric lymph node expansion were measured. WT AOM/DSS mice showed less body weight gain and increased mensenteric lymph node expansion when compared to GPR4 KO AOM/DSS mice. (C) Final body weight change percentage, (D) mensenteric lymph node size, and (E) colon lenghth. Data are presented as the mean ± SEM. Statistical significance was determined using the unpaired t-test between WT AOM/DSS and GPR4 KO AOM/DSS mice. (*P < 0.05, **P < 0.01, ***P < 0.001, and ****P < 0.0001).

### 3.3. GPR4 knockout reduces tumor burden in the CAC mouse model

A significant increase in colon tumor number was observed in WT AOM/DSS mice when compared to GPR4 KO AOM/DSS mice (Fig. 5A-B), suggesting GPR4 promotes CAC development. WT mice had an average tumor number of 7.4 whereas GPR4 KO mice had an average of 5.1, demonstrating a 45.1% increase in tumor number in the WT over GPR4 KO AOM/DSS mice (Fig. 5B). Moreover, the volume of the detected tumors was 115 mm^3^ in WT mice and 55 mm^3^ in GPR4 KO AOM/DSS mice, respectively (Fig. 5C). Histological analyses of the colon tissue sections revealed colon dysplasia (Dys) and adenocarcinoma in situ (AIS) were induced in both WT and GPR4 KO AOM/DSS mice (Fig. 5D-E). Interestingly, the distribution of dysplasia and adenocarcinoma in WT and GPR4 KO AOM/DSS mice showed a 9% decrease in adenocarcinomas in GPR4 KO mice compared to WT (85% vs 94%, respectively) (Fig. 5E). These data suggest absence of GPR4 delays progression from low grade dysplasia to adenocarcinoma in the inflamed colon tissues.

**Fig. 5.**
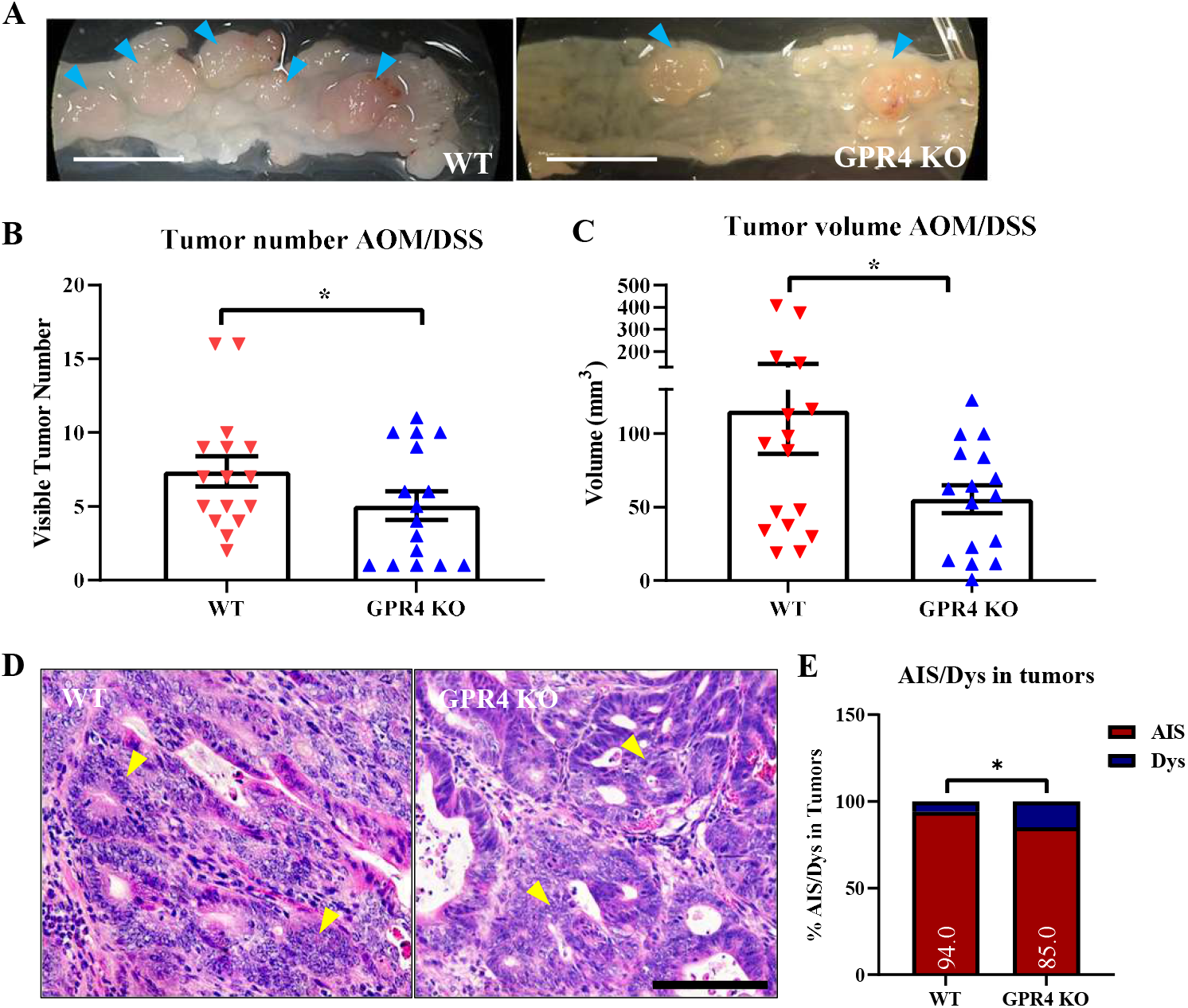
Effects of GPR4 on tumorigenesis in the colitis associated colorectal cancer (CAC) mouse model. Tumor burden was evaluated using clinical observation of tumor number and tumor volume at the endpoint for WT AOM/DSS (n=16) mouse colons compared to GPR4 KO AOM/DSS (n=16). Tumor burden was found to be higher in WT AOM/DSS mouse colons compared to GPR4 KO AOM/DSS. (A) Representative pictures of distal colons bearing tumors (indicated by blue arrowheads), (B) tumor number, (C) tumor volume, (D) histopathological representation of adenocarcinoma in AOM/DSS mouse colons (yellow arrowheads), and (E) distribution of adenocarcinoma in situ (AIS) and low-grade dysplasia (Dys) in the WT AOM/DSS and GPR4 KO AOM/DSS mouse tumors. Data are presented as the mean ± SEM and statistical significance was determined using the unpaired t-test and chi-square test between WT AOM/DSS and GPR4 KO AOM/DSS mice. (*P < 0.05). Scale bar in (A) is 1cm and scale bar in (D) is 100µm.

### 3.4. GPR4 is highly expressed in the tumor blood vessels of AOM/DSS mice

We next characterized the expression profile of GPR4 in the AOM/DSS mouse tumor model. Immunohistochemistry was performed for green fluorescence protein (GFP) which functions as a surrogate marker for GPR4 expression in the GPR4 KO mice. GPR4 KO mice were generated by replacing the GPR4 coding region with an internal ribosome entry site (IRES)-GFP cassette under the control of the endogenous GPR4 gene promoter as previously described [16,19].

GFP signal was detected in GPR4 KO AOM/DSS, but not WT AOM/DSS colon tissues. In line with our previous observations [16], high levels of GFP expression were detected in endothelial cells of tumor blood vessels (Fig. 6A). Furthermore, using a double flourescent stain with GFP and the endothelial marker CD31, we observed that GFP expression was predominately detected in the endothelial cells (ECs) of blood vessels in the tumor tissue of GPR4 KO AOM/DSS colons (Fig. 6B).

**Fig. 6.**
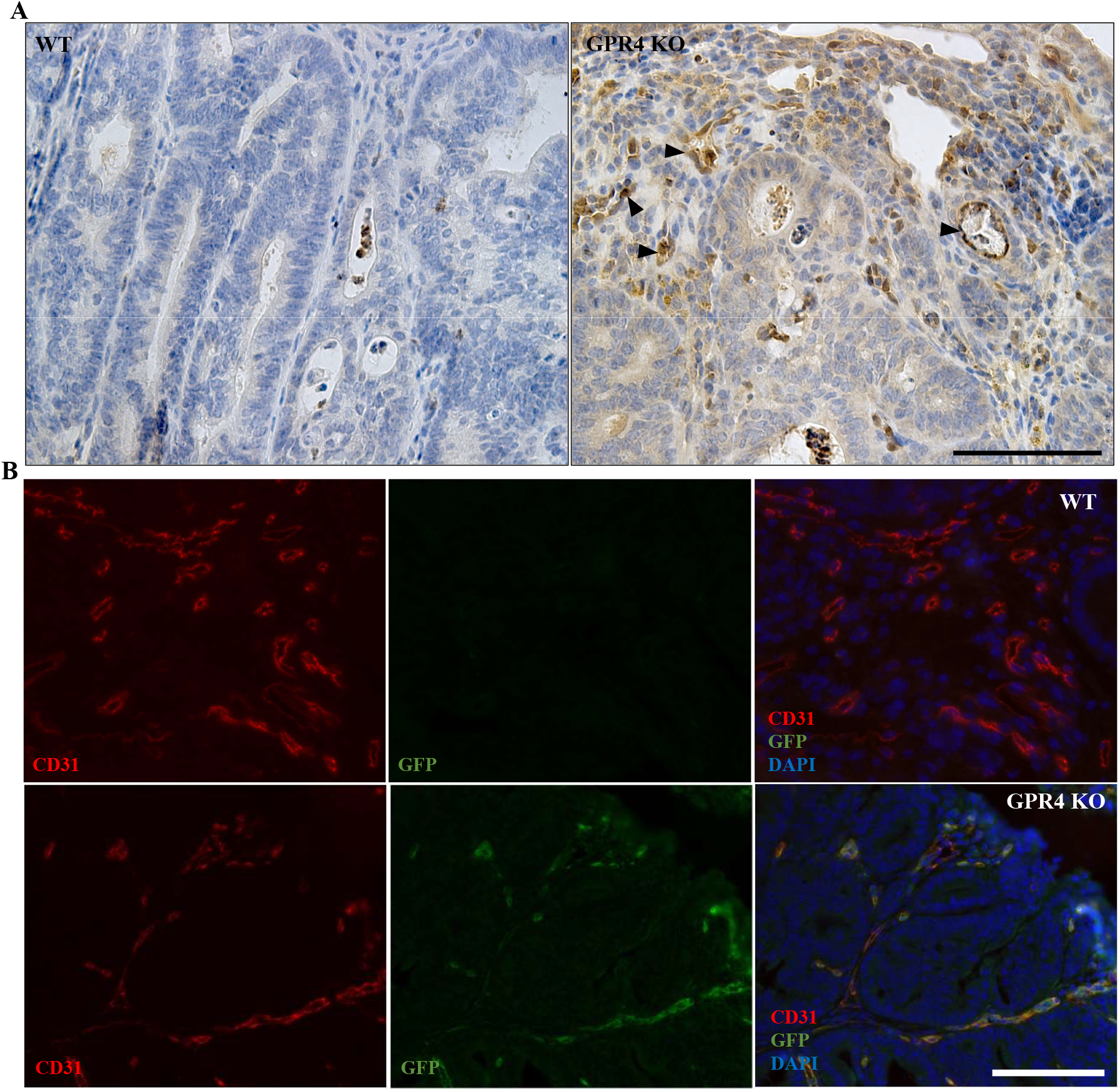
GFP signal and double labeling with CD31 in the colon tumor tissues of the colitis associated colorectal cancer (CAC) mouse model. GFP knock-in under the control of the GPR4 promoter serves as a surrogate marker for endogenous GPR4 expression in GPR4 KO mice. GFP expression could be detected by immunohistochemistry in GPR4 KO mice but not in WT mice. (A) GFP expression (indicated by black arrowheads) in GPR4 KO AOM/DSS tumors versus no expression in WT AOM/DSS tumors, and (B) double labeling of GFP (green) and CD31 (red). Blood vessels show a double positive signal of GFP and CD31 in the GPR4 KO AOM/DSS tumors versus only CD31 positive signal in the WT AOM/DSS tumors. Scale bar is 100µm.

### 3.5. GPR4 deletion decreases angiogenic blood vessel formation in the tumors of AOM/DSS mice

During tumorigenesis, angiogenesis is a fundamental process in the tumor development and progression [41]. We analyzed the blood vessel density in the tumors of WT AOM/DSS and GPR4 KO AOM/DSS using immunohistochemistry. The endothelial marker CD31 was used to identify tumor blood vessels for counting. The deficiency of GPR4 in the GPR4 KO AOM/DSS mice caused a significant reduction in blood vessel density in the tumors of these mice by ∼2.6 fold when compared to WT AOM/DSS (Fig. 7A-B).

**Fig. 7.**
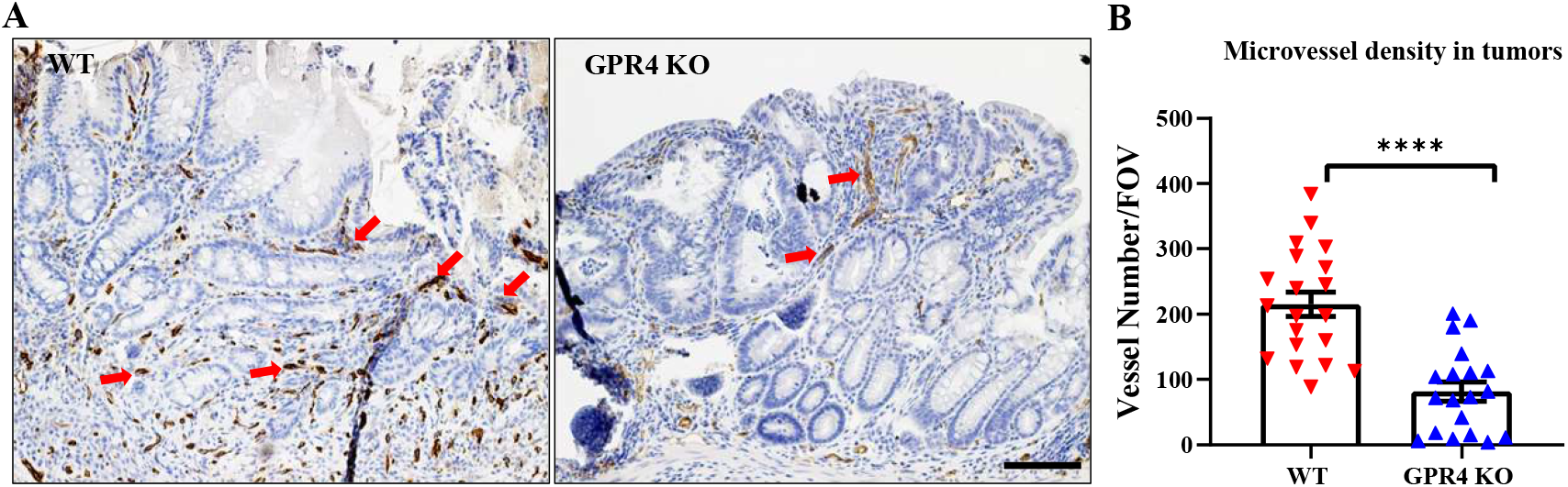
Microvessel density in the tumor tissues of the colitis associated colorectal cancer (CAC) mouse model. Using CD31 immunohistochemistry, blood vessel numbers were assessed in WT AOM/DSS (n=20 tumors) and GPR4 KO AOM/DSS (n=19 tumors). WT AOM/DSS tumors showed increased number of blood vessels compared to GPR4 KO AOM/DSS tumors. (A) Representative pictures of WT and GPR4 KO AOM/DSS tumor CD31+ blood vessels (red arrows), and (B) quantification of average microvessel density per field of view (FOV) in the tumors. ImageJ was used for quantification. Data are presented as the mean ± SEM and statistical significance was determined using the unpaired t-test (****P < 0.0001). Scale bar is 100µm.

### 3.6. GPR4 knockout increases necrosis, cell death and decreases cell proliferation in the tumors of AOM/DSS mice

Tumor necrosis is considered as a common feature of solid tumors which occurs as a consequence of nutrient and oxygen depravation [42]. Areas in the tumors with apparent gaps resulting from transformed crypt loss were identified as necrotic areas, where remenants of nuclear and cytolplasmic debris were visualized with H&E stain (Fig. 8A). The percentage of tumor necrosis area between WT and GPR4 KO AOM/DSS tumors observed to be ∼ 2.4 fold higher in GPR4 KO AOM/DSS compared to GPR4 WT AOM/DSS tumors (Fig. 8A-B). Using immunohistochemistry, we assessed the protein expression of cleaved caspase-3, a marker of cell death, in WT and GPR4 KO AOM/DSS tumors. The expression of cleaved caspase-3 was predominantly detected in the necrotic tumor area in addition to positive epithelial cells scattered within the tumor (Fig. 8C). Next we evaluated Ki67 as a proliferation marker to asses tumor cell proliferation in the WT and GPR4 KO AOM/DSS mice. WT AOM/DSS tumors show a more proliferative phenotype when compared to GPR4 KO AOM/DSS (Fig. 8D). Quantification of Ki67 positive cells revealed a ∼ 2 fold decrease in proliferating tumor cells in GPR4 KO AOM/DSS mouse tumors (Fig. 8E).

**Fig. 8.**
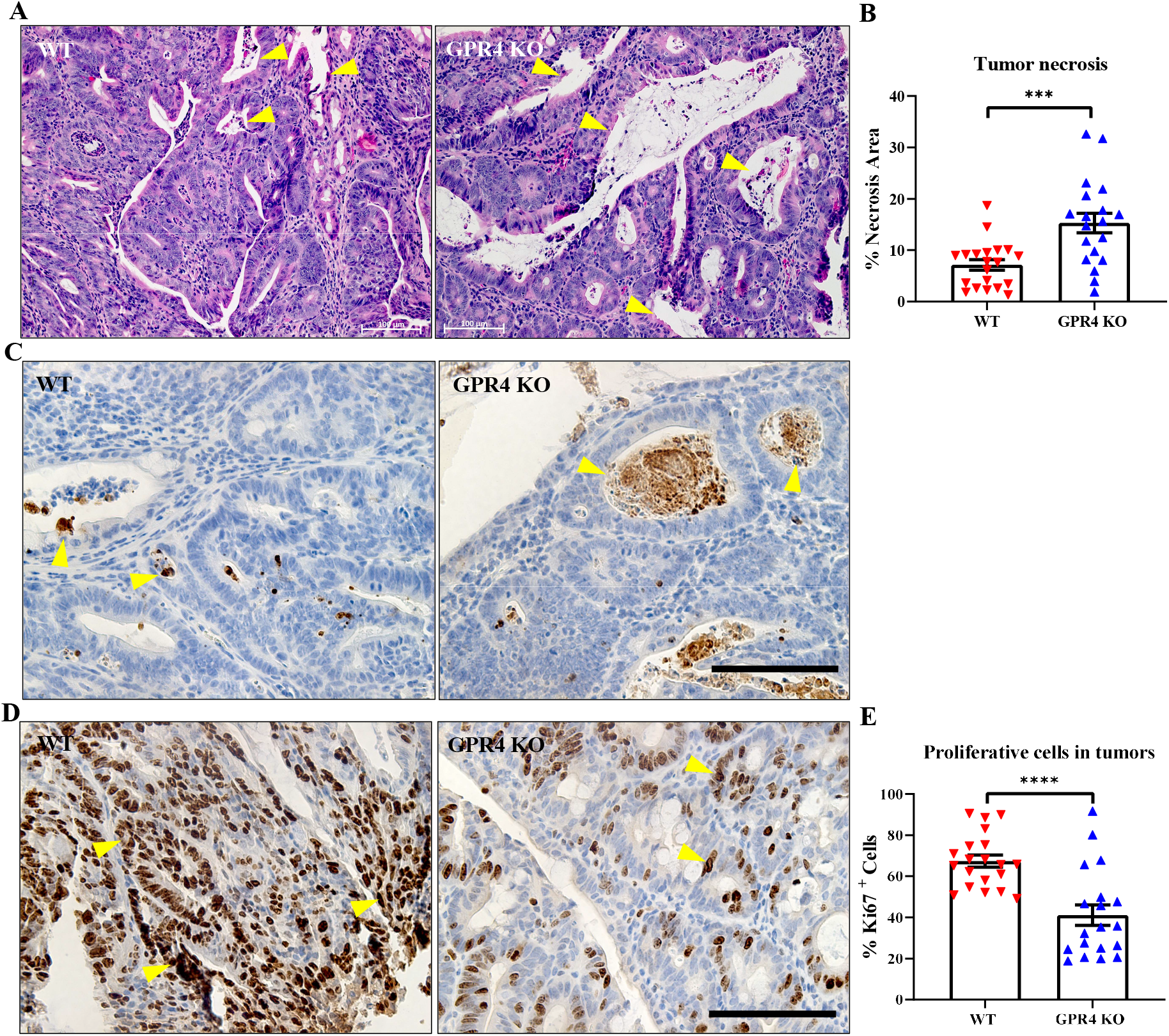
Tumor necrosis, cell death and proliferation in the tumor tissues of the colitis associated colorectal cancer (CAC) mouse model. H&E stain was used to assess necrotic areas in WT AOM/DSS (n=20 tumors) and GPR4 KO AOM/DSS (n=19 tumors) mouse tumors. GPR4 KO AOM/DSS showed higher percent of tumor necrosis when compared with WT AOM/DSS. (A) Representative images of H&E stained tumor tissues showing necrotic areas (yellow arrowheads), and (B) quantification of average percent of tumor necrosis area in WT AOM/DSS and GPR4 AOM/DSS mouse tumors. Cell death was assessed using cleaved caspase-3 immunohistochemistry in the tumor tissues of WT AOM/DSS and GPR4 KO AOM/DSS. (C) Representative images of cleaved caspase-3+ cells (yellow arrowheads) in WT AOM/DSS and GPR4 AOM/DSS mouse tumor tissues. Ki67+ IHC was used to assess cell proliferation in WT AOM/DSS (n=20 tumors) and GPR4 KO AOM/DSS (n=19 tumors) mouse tumors. WT AOM/DSS tumors showed increased percent of proliferative cells when compared with GPR4 KO AOM/DSS tumors. (D) Representative images of Ki67+ (yellow arrowheads) stained tumor tissues for WT AOM/DSS and GPR4 KO AOM/DSS mice, and (E) quantification of average percent of Ki67+ proliferative cells in WT AOM/DSS and GPR4 AOM/DSS mouse colon tumors. Images were quantified using the Fiji software. Data are presented as the mean ± SEM and statistical significance was determined using the unpaired t-test (***P<0.001, ****P<0.0001). Scale bar is 100µm.

## 4. Discussion

In this study we demonstrate that GPR4-driven intestinal inflammation can increase the development of colorectal tumorigenesis in a colitis-associated colorectal cancer mouse model. Our observations are in line with previous reports establishing a proinflammatory role for GPR4 in several systems such as the brain, heart, kidney, lung, bone, skin, and the gastrointestinal tract [16,18,19,20,33,43,44,45,46,47,48,49]. In addition, GPR4 has a protumorigenic role in hepatocellular, head and neck, breast and colorectal cancers [27,28,29].

We and others have previously elucidated the role of GPR4 in potentiating intestinal inflammation by use of both GPR4 antagonists and global GPR4 knockout mice in colitis mouse models [16,18,20,33]. These studies have demonstrated a proinflammatory role for GPR4 in colitis by activation of the intestinal endothelium to facilitate immune cell recruitment and infiltration into the inflamed intestinal tissues [16,18,20,33]. Furthermore, GPR4 mRNA expression levels are significantly increased in the inflamed intestinal tissues of both human IBD patients and in colitis mouse models when compared to normal tissues. These data implicate GPR4 in the pathophysiology of chronic intestinal inflammation [16,18].

Patients with IBD are at a ∼ 2.4 fold higher risk for developing colitis associated colorectal cancer (CAC) due to persistent and unresolved intestinal inflammation [2]. Local tissue acidosis exists in the inflammatory loci and is owing in part to shifts in hypoxia, glycolytic cellular metabolism, production of bacterial metabolic byproducts, and granulocyte respiratory bursts [6,7,8,50]. Increased proton concentrations in the extracellular milieu can activate GPR4 by protonation of histidine residues on the extracellular domain and initiate activation of downstream G protein pathways [51]. We and others have previously demonstrated GPR4 activation in endothelial cells elicits production of inflammatory molecules such as TNF and NF-κB family members, chemokines, cytokines, and adhesion molecules, and functionally mediates leukocyte adhesion to the activated endothelium [16,23,33,52]. In this study, we demonstrated that GPR4 increased intestinal inflammation in the DSS chronic colitis and the AOM/DSS colitis-associated colorectal cancer mouse models (Figs. 1-4). Also, a positive correlation between evelvated GPR4 expression with elevated TNF-α and IFN-γ expression in intestinal tissues of IBD patients was observed (Fig. 3). We have previously shown GPR4 antagonists in the DSS-induced colitis mouse model reduced TNF-α mRNA levels in intestinal tissues when compared to vehicle control [33]. In line with these observations, GPR4 KO mice in the DSS-chronic colitis mouse model had reduced TNF-α expression in the intestinal tissues and less TNF-α-positive infiltrating leukocytes present in the mucosa (Fig. 3). Several studies have linked activation of TNF-α and NF-κB pathways with initiating malignant transformation of the intestinal epithelial cells [53,54]. These data are supported by studies employing genetic and pharmacological blockade of TNF-α which reduced CAC tumor burden in mice [55,56]. In this context GPR4 contributes to chronic inflammation and tumor development which can feedforward in the inflammation-dysplasia-carcinoma axis resulting in CAC development [1].

Interestingly, it has also been reported GPR4 expression is increased in hepatocellular carcinoma, head and neck cancer and CRC tissues compared to normal tissues [27,29,30]. Additionally, hepatocellular and CRC patients with high GPR4 expression showed poor prognosis and decreased survival [29,30]. Furthermore, a significant reduction in breast and CRC tumor growth in the GPR4 knockout mouse and in vitro CRC cell proliferation by GPR4 knockdown, respectively, was observed [28,30]. These observations are in line with our findings where we observe reduced colon tumor burden in GPR4 KO mice compared to WT in the AOM/DSS mouse model (Fig. 5).

Another protumorigenic role for GPR4 is likely to be related to angiogenesis [27,28,29]. GPR4 has been shown to promote angiogenesis by regulating the VEGF pathway in both ischemic tissues and cancer tissues [20,28,47]. One study showed that upregulating GPR4 enhanced angiogenesis of endothelial progenitor cells (EPCs) isolated from chronic artery disease patients through VEGFA/STAT3 activation. The study also showed increased blood flow in a mouse hindlimb ischemia model following GPR4-overexpressing EPC injection [47]. Another study demonstrated a reduced growth of orthotopic breast cancer and colon cancer allografts in GPR4 KO mice due to decreased angiogenesis [28]. Consistently, GPR4 KO mouse tumors showed reductions in tumor angiogenesis when compared to WT tumors in the AOM/DSS mouse model (Fig. 7). We have previously characterized the expression of GPR4 in intestinal tissues of both naïve and the DSS-induced colitis disease state using GPR4 KO mice with a GFP knock-in [16]. In this system, GFP is under the regulation of the endogenous GPR4 promoter and can be used as a surrogate marker for endogenous GPR4 expression. We observed GPR4 is predominately expressed in the vascular endothelial cells of arteries, veins, and microvessels of both the cecum and colon [16]. In this study we employed double immunolabeling of GFP and CD31 to detect GPR4 expression in tumor vascularization (Fig. 6). We found GPR4 is expressed in the vascular endothelial cells of colorectal tumors in AOM/DSS mice and may be involved in tumor vascularization. As shown in previous studies, GPR4 may regulate angiogenesis through the VEGF/VEGFR pathway, STAT3, and the production of proangiogenic factors [20,27,28,47]. Our observations in this study suggest that the decrease in microvessel density in tumors of GPR4 KO AOM/DSS mice is associated with increased tumor cell death and reduced tumor cell proliferation by preventing adequate tumor vascularization (Figs. 7-8). Collectively, genetic deletion of GPR4 likely halts colorectal cancer development by dampening chronic intestinal inflammation and impeding tumor angiogenesis.

The therapeutic benefit for GPR4 antagonism has been described in reducing inflammation, pain, and angiogenesis [22,43,45,48,52,57,58,59]. We and others have shown the therapeutic value for GPR4 antagonists in IBD pre-clinical mouse models [20,33]. Additional benefits for GPR4 antagonism have been reported in tissue ischemia, myocardial infarction, chronic obstructive pulmonary disease (COPD), and osteoarthiritis models, by redcuing proinflammatory molecules such as VCAM-1, E-selectin, IL-17, interferon-γ, TNF-α, IL-1β, IL-6, inducible nitric oxide synthase (iNOS), nitric oxide (NO), cyclooxygenase 2 (COX2), prostaglandin E2 (PGE2), Mucin5AC, matrix metalloprotease (MMP)-9, MMP-12, and NF-κB [22,43,45,48,59]. Based on the research findings, we propose GPR4 inhibition (e.g., via small molecule antagonists, siRNAs, antisense oligos, and antibodies) can be explored as a potential therapeutic approach for colitis treatment and CAC prevention in IBD patients by alleviating chronic intestinal inflammation and inhibiting pathological angiogenesis.

## Acknowledgement

We thank Dr. Owen N. Witte for providing the GPR4 knockout mouse strain.

